# Phylogenomics of *Rhinogobius* gobies reveal northern-southern divergence and trait evolution in China

**DOI:** 10.64898/2025.11.30.691486

**Authors:** Suhan Liu, Qinwen Xue, Yun Hu, Jiantao Hu, Yiling Pan, Yun Bu, Jiliang Wang, Jianhong Xia, Chenhong Li

## Abstract

The goby genus *Rhinogobius* represents a prominent radiation of freshwater fishes in East Asia, with China serving as its evolutionary hotspot harboring over 39 endemic species. Despite their ecological dominance in riverine ecosystems, phylogenetic relationships among mainland Chinese *Rhinogobius* species remain poorly resolved due to limited sampling and morphological conservatism, particularly in meristic traits traditionally used for classification. To address these knowledge gaps, we employed a nuclear gene enrichment approach to target 4,434 single-copy nuclear loci across 26 species spanning all major river basins in eastern and southern China. Our phylogenomic analyses based on sequence of 2,137 cleaned-up loci (477,822 bp) recovered a robust topology dividing *Rhinogobius* into three primary lineages: an early-diverging of *R. similis* sister to two major clades, the northern vs. southern clades exhibiting strong geographic structuring. Fossil-calibrated molecular dating placed the crown age of *Rhinogobius* at ∼9.33 million years ago (Ma) during the late Miocene, with a pivotal north-south divergence at ∼5.96 Ma coinciding with major paleo-drainage reorganizations in East Asia. Ancestral state reconstruction revealed that the southern clade uniquely evolved unstable first dorsal fin formula and near-universal presence of predorsal scales. By integrating the most extensive geographic and taxonomic sampling to date with phylogenomic data, this study establishes: (1) a revised evolutionary timeline for *Rhinogobius* diversification linked to East Asian geomorphological history, (2) diagnostic morphological characters for major clades, and (3) a robust phylogenetic framework to guide future taxonomic revisions and conservation efforts for this ecologically significant group.

**Highlights:**

- Phylogenomics resolves north/south split in Chinese *Rhinogobius* gobies.
- Fossil-dated 9.3 MYA origin of *Rhinogobius* and rapid divergence at 5.96 MYA.
- Southern lineages exhibit more predorsal scales and unstable first dorsal fin formula.
- First genomic backbone for species revision and adaptation study.

## 1. Introduction

The genus *Rhinogobius* Gill, 1859 (family Oxudercidae) comprises the most speciose and geographically diversified freshwater gobiid radiation in East and Southeast Asia, occupying river systems from Vietnam to Japan (Yamasaki et al., 2015). China represents the evolutionary center of this genus, containing at least 42 species (39 endemic) distributed across mainland China, Taiwan, and Hainan Island (Wu, 2008). These ecologically versatile fishes demonstrate extraordinary adaptive traits, including specialized egg morphology (Yamasaki and Tachihara, 2015; Katoh and Nishida, 2008) and complex reproductive behaviors (Kano et al., 2012), making them exceptional models for studying freshwater adaptive radiation. Despite their ecological dominance and economic value, particularly *R. similis* in fisheries and aquaculture and other species for ornamental trade, their phylogenetic architecture remains poorly understood. Current knowledge derives primarily from studies of Japanese populations (Yamasaki et al., 2015), leaving Chinese remarkable species richness genetically unexplored.

Taxonomic confusion plagues *Rhinogobius* due to conserved morphology and rapid speciation. Discrepancies in species counts persist between authoritative sources: 83 valid species in Catalog of Fishes (Fricke et al., 2019) versus 65 accepted from 108 nominal taxa in FishBase (Froese and Pauly, 2022; Kang et al., 2022). Molecular studies have uncovered extensive synonymy, including *R. similis*-*R. giurinus* (Suzuki et al., 2015) and unrecognized diversity within the *R. brunneus* complex (Akihito et al., 2002, 2013). While species descriptions continue accelerating (Xia et al., 2018; Chen et al., 2022, 2024a, 2024b; Li et al., 2024), most taxa exhibit overlapping meristic ranges (Maeda et al., 2024) and only subtle chromatic differences. This morphological conservatism challenges traditional taxonomy that relies on vertebral counts, scale arrangements, and fin ray formulae, characters increasingly shown to lack diagnostic resolution among sister species.

The phylogenetic relationships within *Rhinogobius* remain controversial, with most studies to date relying exclusively on mitochondrial markers. In the first comparative mitogenomic analysis of five species (*Rhinogobius brunneus*, *R. leavelli*, *R. cliffordpopei*, mainland Chinese *R. similis*, and Taiwanese endemic *R. rubromaculatus*), Zhang and Shen (2019) identified *R. similis* as the basal lineage, a finding subsequently supported by multiple mitochondrial DNA studies (Han et al., 2025; Song et al., 2022; Zhang and Shen, 2019; Tuncharoen et al., 2025). However, Tan et al. (2020) recovered an alternative topology with *R. giurinus* (synonymy of *R. similis*) and *R. leavelli* as sister species. The most comprehensive mitogenomic study to date (Jia et al., 2025) analyzed 16 species, resolving two primary clades: one containing *R. wuyanlingensis*, *R. filamentosus*, *R. zhoui*, and *R. duospilus*, and the other comprising *R. brunneus*, *R. leavelli*, *R. davidi*, *R. cliffordpopei*, *R. wuyiensis*, *R. lentiginis*, and *R. niger*. Notably, this study excluded *R. similis* but placed *R. formosanus* at the base of the genus, contrasting with Han et al. (2025) who grouped *R. formosanus* with *R. similis* in the earliest diverging lineage. These conflicting topologies likely stem from two fundamental limitations of mitochondrial data: their inheritance as a single linkage unit in vertebrates, which conflates gene and species histories, and insufficient taxonomic sampling that fails to capture full diversity of the genus. Consequently, while mitochondrial markers have provided initial insights into *Rhinogobius* phylogeny, their inability to consistently resolve relationships underscores the need for nuclear genomic data to reconstruct robust species trees.

Here, we collected 54 specimens representing 26 *Rhinogobius* species from major river systems across southern and eastern China. Through targeted sequence capture of 4,434 nuclear protein-coding loci, we generated phylogenomic datasets for each individual. Additionally, all voucher specimens were re-examined for 8 meristic characters, supplemented by published morphological data (Wu, 2008; Zhou et al., 2024). This study addresses three primary objectives: (1) reconstruction of the first species-level nuclear phylogeny for *Rhinogobius* using genome-level data; (2) estimation of divergence times and biogeographic history of mainland Chinese lineages; and (3) evaluation of traditional diagnostic characters within a phylogenetic framework. Our integrated phylogenomic-morphological approach resolves long-standing systematic uncertainties while establishing a foundation for future taxonomic revisions.

## 2. Materials and methods

### 2.1. Sampling and DNA extraction

Fifty-four specimens representing 26 *Rhinogobius* species were collected from major river systems in southern and eastern China, including the Yangtze, Qiantang, Oujiang, Hanjiang, Rongjiang, Red, and Liujiang river basins, as well as the isolated drainages of Hainan Island (Fig. 1; Table 1). There are 16 described species from the 26 species and 10 undescribed species (details of these can be found in Supplementary Table S1). Fin clips or muscle tissue samples were preserved in 95% ethanol immediately after collection. Two *Tridentiger* species (*T. barbatus* and *T. bifasciatus*), along with *Pterapogon kauderni* and *Sphaeramia orbicularis* representing the Apogonidae, *Kurtus gulliveri* representing the Kurtidae, *Odontobutis potamophilus* representing the Odontobutidae, *Butis koilomatodon*, *Kribia nana*, and *Oxyeleotris marmoratus* representing the Butidae, as well as *Valenciennea puellaris* representing the Gobiidae, were selected as outgroup taxa based on recent gobioid phylogenies (Kuang et al., 2013). All specimens were assigned unique voucher numbers (Supplementary Table S1) and deposited in the Fish Collection of Shanghai Ocean University (SOU), China, (contact person: Dr Ya Zhang). Genomic DNA was extracted from ethanol-preserved tissues using the Ezup Column Animal Genomic DNA Purification Kit (Sangon Biotech, Shanghai, China). DNA concentration and quality were assessed using a NanoDrop 3300 Fluorospectrometer (Thermo Fisher Scientific, Wilmington, DE, USA) and verified by 1% agarose gel electrophoresis. Only samples with A260/A280 ratios between 1.8-2.0 and showing minimal degradation were used for downstream analyses.

**Fig. 1.**
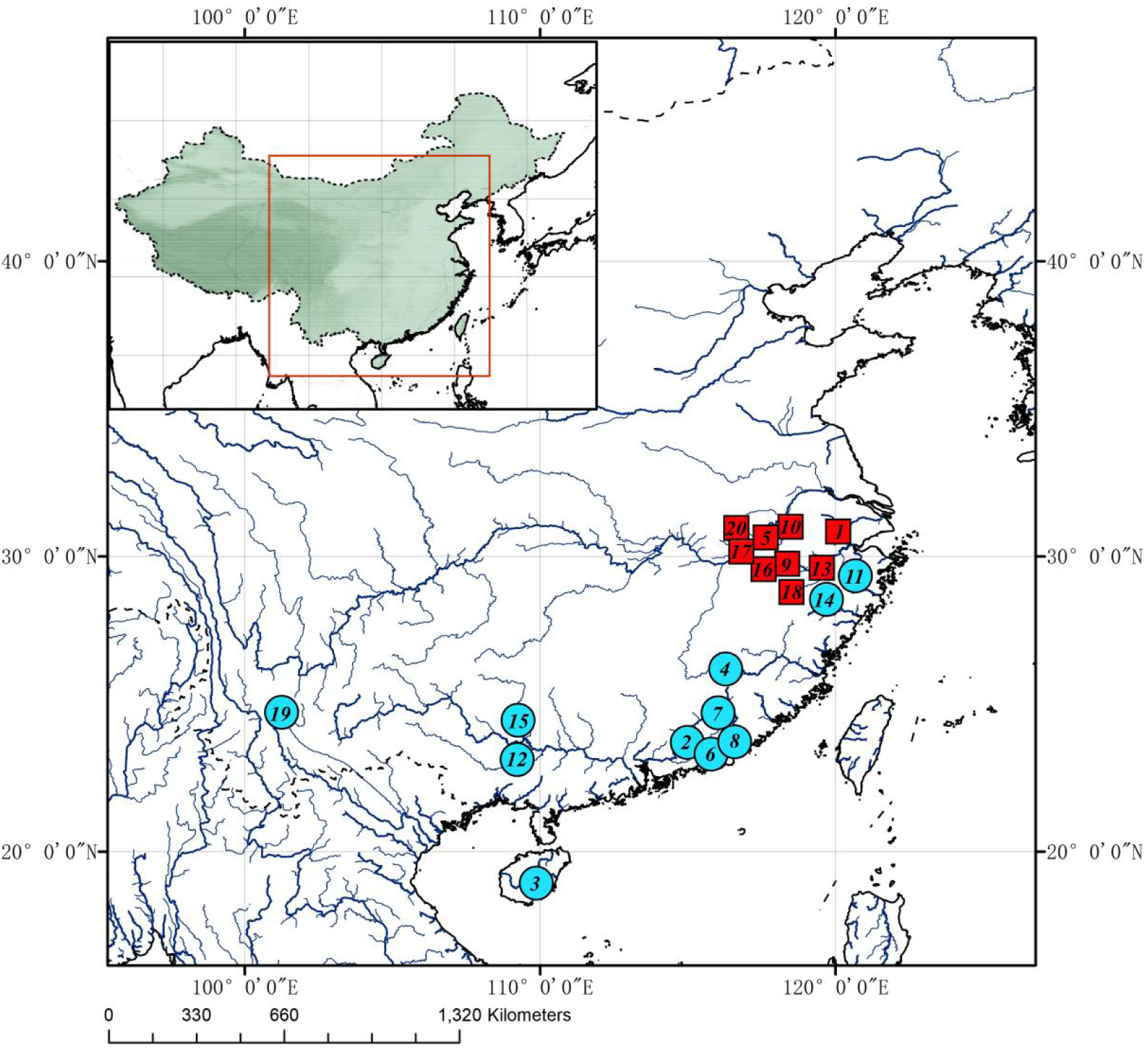
Sampling locations of *Rhinogobius* across major river basins in China. Collection sites are indicated by colored symbols corresponding to phylogenetic clades identified in this study (see Fig. 2). The map highlights rivers with a discharge >40 m³/s, including eight major drainage systems sampled: Yangtze, Qiantang -belonged to northern basin affiliation, sampling sites are marked with red square; Oujiang, Hanjiang, Rongjiang, Red, and Liujiang Rivers, plus the isolated river systems of Hainan Island -belonged to southern basin affiliation, sampling sites are marked with blue circle. Inset shows regional context within East Asia. Base map generated using ArcGIS v10.6 (ESRI, 2022) with hydrographic data from HydroSHEDS (Lehner & Grill, 2013)

**Table 1.** List of samples used in this study. Locality numbers correspond to those in Figure 1.

| Species | specimen ID | River/River system | Locality no. |
| --- | --- | --- | --- |
| *R. sp. ORAZJ* | 2228_3, 2229_4 | Anji County, Yangtze River Basin | 1 |
| *R. multimaculatus* | 2530, 2531 | Anji County, Yangtze River Basin | 1 |
| *R. zhoui* | 1902_1,2, 1904_1 | Haifeng County, Mt.Lianhua | 2 |
| *R. sangenloensis* | 2222_2,3 | Hainan | 3 |
| *R. changtinensis* | 2495_1 | Hanjiang River | 4 |
| *R. sp. LLVC27* | 090801, 090803 | Huangshan County, Yangtze Basin | 5 |
| *R. sp. XIAO* | 1901_1,5,8 | Jiexi County, Rongjiang River | 6 |
| *R. sp. YLGD* | 1903_2,3,4,6 | Jieyang City, Hanjiang River | 7 |
| *R. duospilus* | 1908_2,4 | Jieyang City, Rongjiang River | 8 |
| *R. cf. lentiginis* | 2226_2,3 | Kaihua County, Qiantang Basin | 9 |
| *R. niger* | 2224_1,2 | Lin'an City, Yangtze River Basin | 10 |
| *R. genanematus* | 2215_1,4 | Lingjiang River | 11 |
| *R. lentiginis* | 1328_1,6 | Lingjiang River | 11 |
| *R. cliffordpopei* | 1147_2,9 | Liujiang River | 12 |
| *R. leavelli* | 2461_2,3 201803012 | Longyou County, Qiantang Basin | 13 |
| *R. wuyanlingensis* | 1850_1,4 | OuJiang | 14 |
| *R. filamentosus* | 2216_1,2,3 | Perl River Basin | 15 |
| *R. davidi* | 2227_1,4 | Qimen County, Qiantang Basin | 16 |
| *R. sp. LLAH* | 2231_4 | Qimen County, Yangtze River Basin | 17 |
| *R. sp. 2 OSVSAH* | 2232_3,4 | Qimen County, Yangtze River Basin | 17 |
| *R. cf. wuyiensis* | 2230_2 | Qimen County, Yangtze River Basin | 17 |
| *R. sp. P18L33AH* | 2233_3 | Qimen County, Yangtze River Basin | 17 |
| *R. maxillivirgatus* | 2016100002, 0095 | Qimen County, Yangtze River Basin | 17 |
| *R. wuyiensis* | 1327_1,2 | Quzhou City, Qiantang River Basin | 18 |
| *R. honghensis* | 2220_3,4,5 | Red River Basin(Yunnan) | 19 |
| *R. sp. CBBSAH* | 2225_2 | Shitai County, Yangtze River Basin | 20 |

### 2.2. Library preparation and targeted sequencing

library preparation and target enrichment procedures were conducted on the basis of established protocols (Meyer and Kircher, 2010) with adaptations for divergent species (Li et al., 2013). Initially, 50 μl of purified DNA was fragmented to ∼250 bp with a SCIENTZ18-A Ultrasonic DNA Interrupter (Scientz, Ningbo, China), employing a protocol consisting of 22 cycles of ultrasonication (20 seconds on, 20 seconds off) at 4°C with 300W processing power. Library construction began with 300 ng of sheared DNA per sample. A set of biotinylated RNA baits (120 bp length, 2× tiling density) was synthesized by MYcroarray (Ann Arbor, Michigan, USA; Catalog #150901-Li-Goby), designed based on 4,434 single-copy nuclear exon markers previously established for phylogenetic studies of gobies (Kuang et al., 2018; Jiang et al., 2019; Hu et al., 2023). Two successive rounds of hybridization capture were performed for each library to optimize target enrichment efficiency. All libraries were uniquely identified with 8 bp dual-index combinations on P5 and P7 adapters to enable multiplexing. Following quality control, enriched libraries were pooled in equimolar concentrations and subjected to 150 bp paired-end sequencing on an Illumina HiSeq X Ten platform (Illumina, San Diego, CA, USA) through Novogene Technology (Beijing, China).

### 2.3. Data processing and assembly

Raw sequence reads were demultiplexed according to the 8 bp sample-specific indices. Subsequently, Cutadapt v1.2.1 (Martin, 2011) and Trim_Galore v0.6.7 (Krueger, 2021) were employed to remove adapter sequences and low-quality bases (Q < 20) from demultiplexed reads by default parameters. Sequence assembly followed Yuan et al. (2019) using a custom Perl pipeline to acquire locus-specific contigs from trimmed reads. The pipeline identified paralogous sequences by aligning putative targets to the reference genome, with sequences failing to match target regions being excluded as probable paralogs. Loci with low enrichment efficiency (< 80% taxonomic coverage and <50% sample coverage) were removed from the final assembly results. The filtered outputs loci were initially aligned in batch using Clustal Omega (Sievers et al., 2011) based on codons, then back-translated to nucleotide alignments while preserving codon structure. The alignment dataset was further refined by excluding loci showing excessive pairwise distances that could not be manually corrected.

### 2.4. Phylogenetic analyses

Aligned sequence were concatenated and subjected to Maximum likelihood (ML) trees reconstruction using RAxML v8.2.8 (Stamatakis, 2014) under the GTR+Γ model with 1,000 bootstrap replicates(Kalyaanamoorthy et al., 2017). Both codon-position partitioning and an optimized scheme selected by PartitionFinder v2.1.1 (Lanfear et al., 2017) were evaluated. The partitioning optimization, initiated from codon-based data blocks, employed the rcluster algorithm under GTR+Γ and selected the best-fit scheme using Bayesian Information Criterion (BIC).

### 2.5. Divergence time estimation

Divergence time estimation was conducted basing on clock-like loci selected from aligned loci in section 2.3 but with 100% of taxonomic coverage and containing at least 90% individuals (n > 58) for all 36 analyzed species (26 ingroup and 10 outgroup taxa). The most clock-like loci were identified using a Perl script developed by Yuan et al. (2019), which rank loci by comparing log likelihood scores between strict-clock and relaxed-clock models for individual gene trees. After excluding loci rejected for molecular clock (p < 0.05), 612 loci were used for divergence time estimation.

Subsequently, these loci were converted into the .nex format and configured in BEAUti v2.7.8 for Bayesian dating analyses in BEAST v2.7.8 (Bouckaert et al., 2019; Drummond et al., 2007) with following parameters: codon-partitioned data (positions 1+2+3) with linked clock models and trees but unlinked site models. The substitution model was GTR+Γ with 4 gamma categories and 1.0 substitution rate for all partitions. Clock model was relaxed clock log normal with 1.0 mean clock rate and tree model was birth-death. To prevent potential convergence failures associated with random starting trees in multiple BEAST runs, the rooted RAxML tree topology obtained from previous concatenated analysis (Section 2.4) was employed as the foundation and subsequently calibrated with fossil dates (listed in the following paragraph) using the penalized likelihood method implemented in r8s. The resulting time-calibrated Newick-formatted tree served as the starting topology for all BEAST analyses. In addition, four parameters: Birth Death Subtree Slide, Narrow, Wide and Wilson-Balding were disabled to ensure tree topology is fixed throughout each BEAST runs. Two independent MCMC runs of 60 million generations each were performed, sampling parameters every 1,000 generations. Convergence was assessed in Tracer v1.7.2 (Rambaut et al., 2018), requiring effective sample sizes > 200 for all parameters. The resulting maximum-clade credibility (MCC) tree with means and 95% highest posterior density of divergence times was generated with TreeAnnotator v2.7.8 after discarding 20% burn-in using LogCombiner v2.7.8 and visualized in FigTree v1.4.4.

Four nodes were constrained using well-documented fossil calibrations within Gobiiformes. The first calibration point established a hard minimum age of 47.8 Ma for the crown group of all gobiiformes species, based on †Carlomonnius quasigobius (Bannikov & Carnevale, 2016), the oldest known gobioid fossil from the Lower Eocene of northern Italy (47.8–56.0 Ma) along with a soft maximum age of 122 Ma (95% CI) derived from mitochondrial-based estimates of the gobiiformes root age (Chakrabarty et al., 2012). This node was modeled with a lognormal prior (mean = 1.9, SD = 0.5, offset = 47.8). The second fossil calibration established age constraints for the most recent common ancestor of Oxudercidae (*Tridentiger* and *Rhinogobius*), with a hard minimum bound at 17.2 Ma based on the oldest skeleton-based member of the Oxudercidae from the Lower Miocene of Turkey with the geological age estimated between 17.2 and 19.5 Ma. (Moritz et al., 2024) and a 95 % soft maximum bound at 56 Ma based on maximum geographical age of †*Carlomonnius quasigobius* (Lognormal prior settings: mean = 2.343, SD = 0.8, offset = 17.2). Fossil for setting hard minimum bound is †*Simpsonigobius nerimanae* Moritz, Elena & Bettina, 2024, which was placed under the Oxudercidae by sharing two phylogenetically informative morphological characters with the family: (1) premaxilla with a weak or absent postmaxillary process and (2) an anal fin with one ray more than in the second dorsal fin (Moritz et al., 2024). For the *Rhinogobius* crown group, a hard minimum of 0.2 Ma was assigned based on Middle Pleistocene fossils from Japan (1.1–0.2 Ma) (Yabumoto, 1987). Though species-level identification remains uncertain currently, generic assignment to the *Rhinogobius* is supported by following meristic traits: (1) 10 abdominal + 16 caudal vertebrae; (2) first dorsal fin with 6 spines, second dorsal fin with 1 spine + 8 soft rays; (3) anal fin with 1 spine + 8 soft rays. The soft maximum for this node (19.5 Ma) aligns with †*S. nerimanae*’s maximum age (Lognormal prior settings: mean = 1.62, SD = 0.8, offset = 0.2). An additional calibration point was applied to constrain the divergence between *Kribia* and *Butis* (hard minimum: 23.03 Ma, based on †*Lepidocottus aries*; soft maximum: 56 Ma, based on †*C. quasigobius*, lognormal prior settings: mean = 2.18, SD = 0.8, offset = 23.03) to facilitate calibration in r8s.

## 3. Results and Discussion

### 3.1. Phylogenetic relationships and divergence patterns

Final analyses included 2,137 loci (477,822 bp) with low enrichment efficiency (< 80% taxonomic coverage and <50% sample coverage) were removed. The concatenated nuclear phylogenetic analysis strongly supports the monophyly of *Rhinogobius* (bootstrap = 100%, Fig. 2, Supplementary Fig.1), with *R. similis* branched first and the other sampled species forming two well-supported clades exhibiting distinct biogeographic patterns. Clade S (bootstrap = 100%) comprises eleven southern Chinese species: *Rhinogobius* sp. XIAO, *R. zhoui*, *R. filamentosus*, *R. wuyanlingensis*, *R. genanematus*, *R. maxillivirgatus*, *R. changtingensis*, *R. duospilus*, *R. honghensis*, *Rhinogobius* sp. YLGD, and *R. sangenloensis*. Clade N (bootstrap = 100%) contains fifteen northern Chinese taxa including *R. lentiginis*, *R.* cf. *lentiginis*, *Rhinogobius* sp. CBBSAH, *Rhinogobius* sp. LLAH, *Rhinogobius* sp. 2 OSVSAH, *R. wuyiensis*, *R. cf. wuyiensis*, *Rhinogobius* sp. LLVC27, *R. niger*, *Rhinogobius* sp. P18L33AH, *R. leavelli*, *R. cliffordpopei*, *R. davidi*, *Rhinogobius* sp. ORAZJ, and *R. multimaculatus*.

**Fig. 2.**
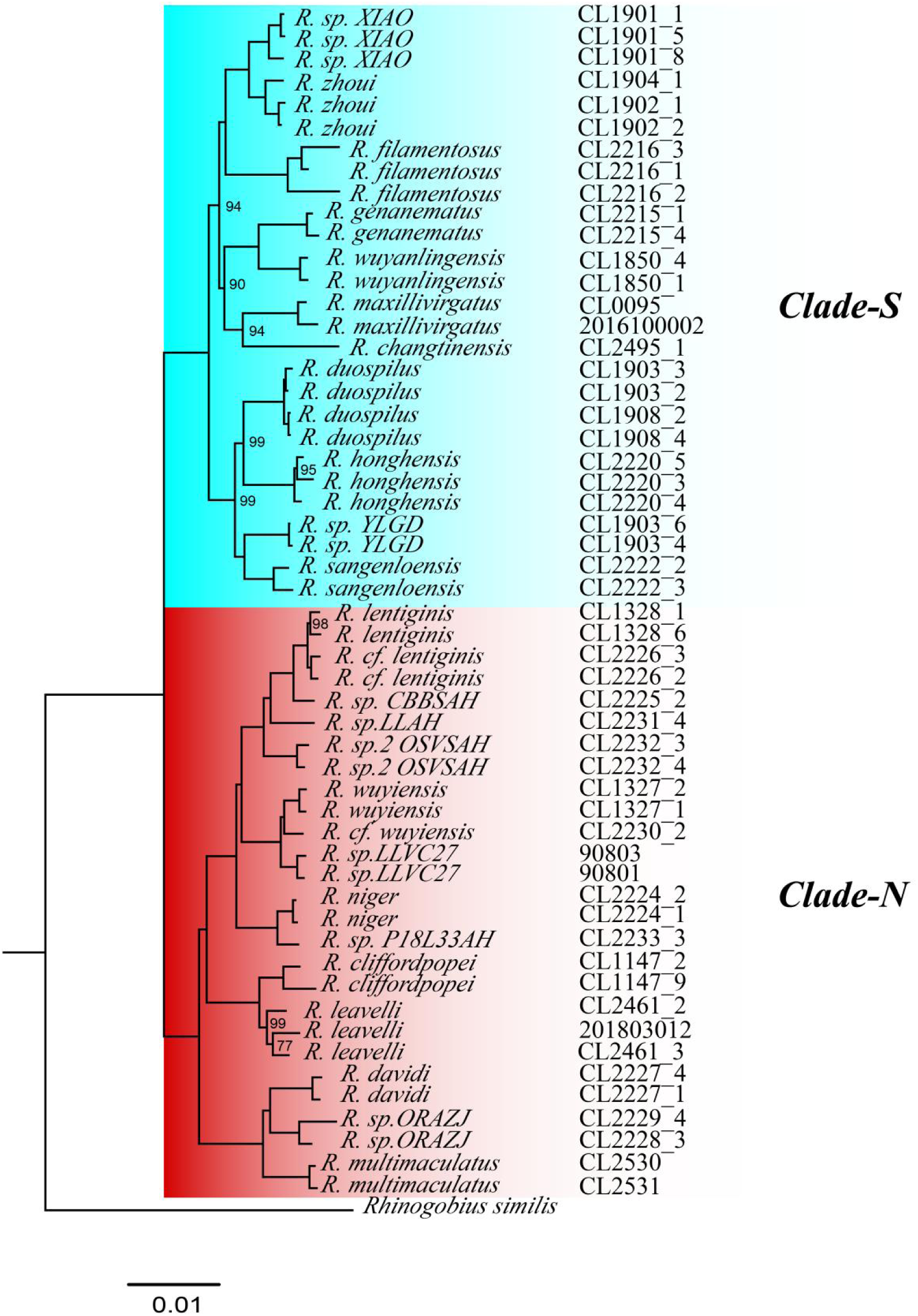
Maximum likelihood phylogeny of *Rhinogobius* based on concatenated analysis of 2,137 nuclear protein-coding loci, reconstructed using RAxML under the GTR+Γ model. Nodes with bootstrap support values <100% are labeled (100% support omitted for clarity). Color gradients at branch tips reflect sampling localities as shown in Fig. 1.

While the major clade structure aligns with Jia et al. (2025), we recovered distinct interspecific relationships within these clades. Notably, *R. zhoui* shows stronger phylogenetic affinity to *R. filamentosus* than to *R. duospilus* (bootstrap = 93%), contrasting with previous mitochondrial-based topologies. The amphidromous *R. similis* forms a distinct early-diverging lineage sister to all other *Rhinogobius* species (bootstrap = 100%), consistent with most mitogenomic studies (Han et al., 2025; Song et al., 2022; Zhang and Shen, 2019). This phylogenetic pattern suggests that the radiation of inland freshwater *Rhinogobius* species in China likely originated from an amphidromous ancestor, supporting a marine-to-freshwater transition in the evolutionary history of this group.

### 3.2. Divergence time and biogeographic pattern

BEAST analysis estimates the origin of *Rhinogobius* in the late Miocene at 9.33 Ma (95% HPD: 6.37 Ma-12.70 Ma), followed by a major divergence at 5.96 Ma (95% HPD: 4.22 Ma-7.92 Ma) that split the genus into the two primary clades identified in the phylogenetic analyses (Fig. 2). This key diversification event predates the Early Pliocene.

The sampled specimens encompass major river basins of eastern and southern China (Fig. 1), which we classified into northern (Yangtze and Qiantang) and southern basins (Lingjiang, Hanjiang, Oujiang, Rongjiang, Pearl, and Liujiang Rivers plus Hainan Island’s drainages) based on geographic position and drainage patterns. Phylogeographic assessment shows a strong but imperfect correlation between clade membership and basin affiliation: while Clade S predominates in southern basins and Clade N in northern basins, with several notable exceptions (Fig. 3A & B).

**Fig. 3.**
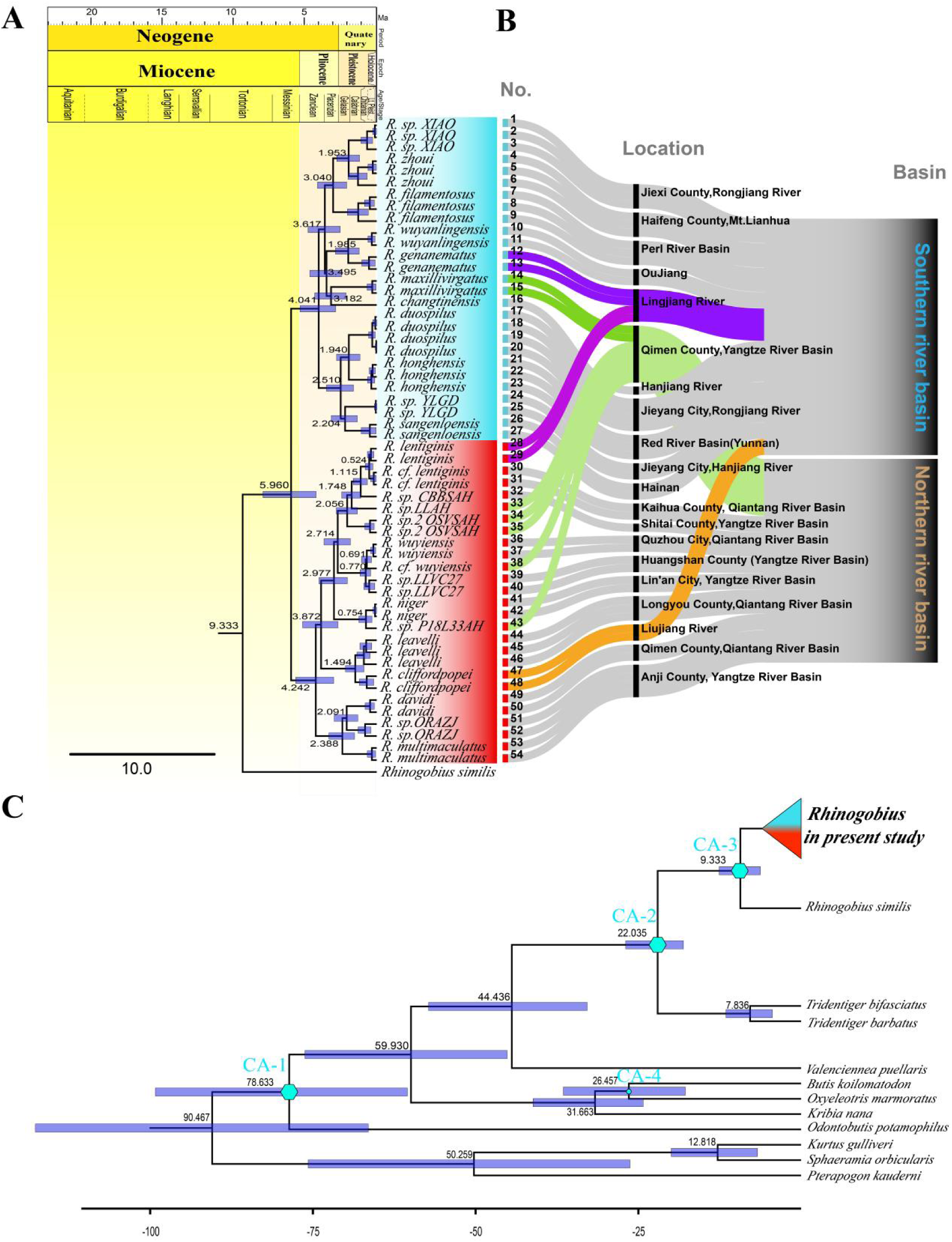
Divergence history and biogeographic relationships of *Rhinogobius* lineages. (A) Time-calibrated species tree of *Rhinogobius* reconstructed using BEAST v2.7.8 (calibration points and outgroups see Figure 3C). Node ages are shown with 95% highest posterior density intervals (blue bars). (B) Alluvial plot illustrating specimen-clade-geography relationships, showing concordant (gray) and discordant (colored) patterns between phylogenetic clades (Clade S [blue] vs. Clade N [red]) and basin affiliations (Southern vs. Northern). (C) Time-calibrated tree showing outgroup of *Rhinogobius*. Calibration point indicated by blue hexagon: CA-1, 47.8–56.0 Ma, *Carlomonnius quasigobius* fossil; CA-2, 17.2-19.5 Ma, *Simpsonigobius nerimanae* fossil; CA-3 0.2-0.8 Ma *Rhinogobius* sp. fossil; CA-4, 23.03-56 Ma, *Lepidocottus aries* fossil.

The genus *Rhinogobius* exhibits a distinct life history polymorphism. Some species display amphidromous traits, while others are entirely landlocked. Drainage systems have played a significant role in driving population divergence among landlocked *Rhinogobius* lineages. Notably, gene flow can be significantly restricted even between adjacent streams less than 5 km apart (Wu et al., 2016). In this research however, two landlocked species (*R. maxillivirgatus* from the Yangtze, northen basins and *R. lentiginis* from the Lingjiang, southen basins) exhibit discordant placements in the phylogenetic clade, suggesting historical gene flow probably mediated by paleo-river reorganization. Conversely, amphidromous species typically exhibit strong dispersal capacity in inland freshwater systems, while showing minimal genetic differentiation among populations (Ju et al., 2016). In this research, the *R*. *cliffordpopei*, also is found in the Liujiang Basin outside our main sampling area, likely represents a more recent range expansion through dispersal. These patterns highlight how both historical river connections and contemporary dispersal capacity have shaped the current distribution of *Rhinogobius* lineages.

### 3.3 Systematic significance of morphological traits and taxonomic implications

#### 3.3.1 Meristic variability challenges morphology-based diagnosis in Rhinogobius

Traditional morphology-based taxonomy of *Rhinogobius* fish relies heavily on meristic characters including vertebral counts, scale patterns (predorsal/longitudinal/transversal), and fin ray numbers. However, our comprehensive analysis of the 26 species (Table 2, Fig. 4) reveals substantial intraspecific variation and interspecific overlap in these traits. Our sample set included numerous topotypic specimens collected from type localities. During morphological examination, we frequently observed discrepancies between morphometric data of these specimens (Table 2) and literature records, providing evidence for intraspecific variation. For example, (1) *R. zhoui*: restricted to the Lianhua Mountain region, our pre-dorsal scale (pre-D) counts (8-9) fell outside the original reported range (10-12; Li & Zhong 2009), as did transverse scale row (TR) counts (7-8 vs 8-9); (2) *R. genanematus*: specimens from the Lingjiang River exhibited fewer vertebrae (25) than documented (27; Chen & Fang 2006) despite matching other characters (Zhong & Tzeng 1998); (3) *R. niger*: TR counts (9-10) were lower than the reported range (10-12; Huang et al. 2016); (4) *R. leavelli*: some specimens showed reduced pre-D counts (5-9) compared to records in *The Gobiidae of China* (Zhou et al., 2024); (5) *R. davidi*: Some specimens displayed fewer vertebrae (27-28 vs 28) and TR rows (7-9 vs 11-12; Chen & Miller 1998).

**Fig. 4.**
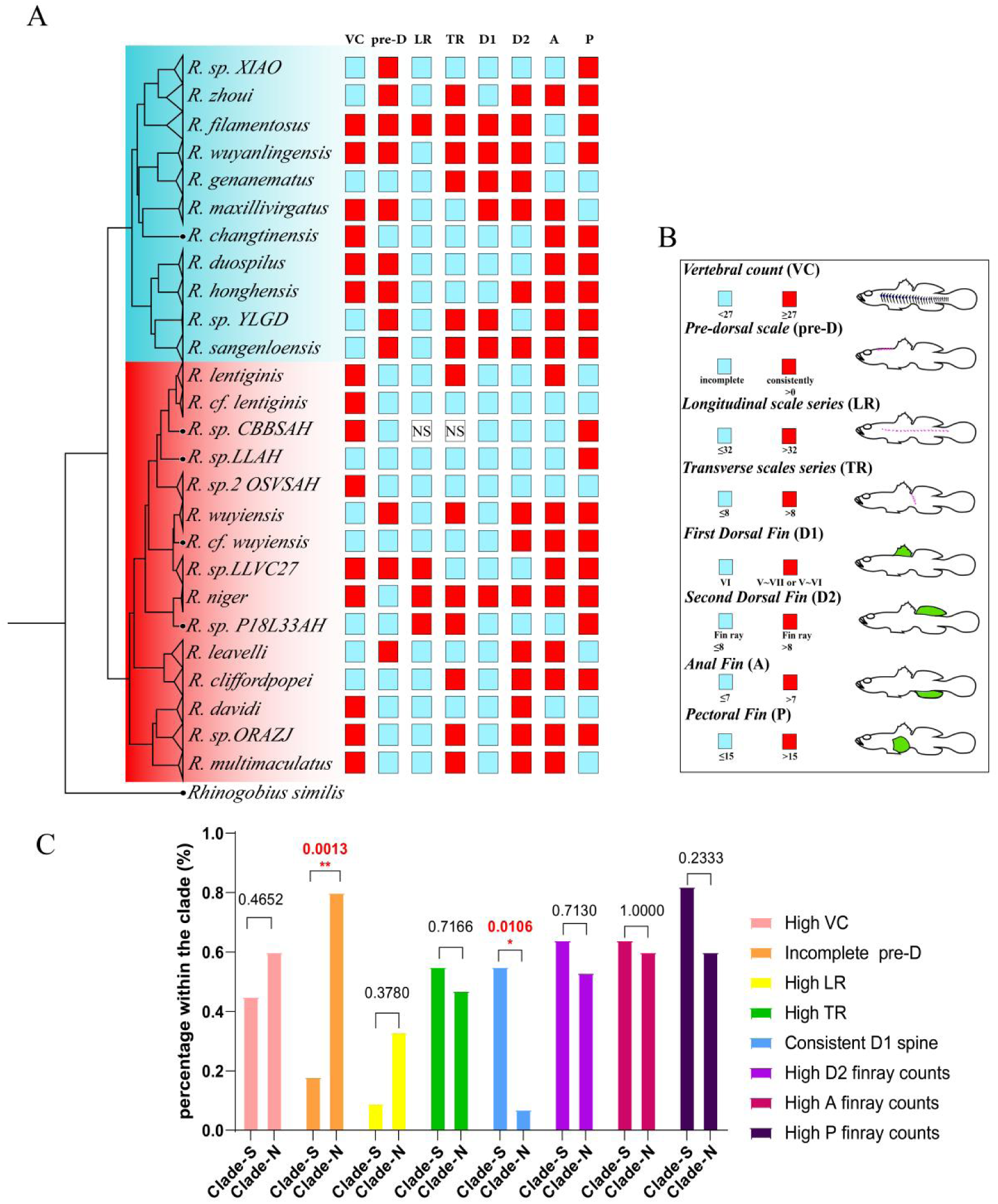
Morphological trait evolution in *Rhinogobius*. (A) Phylogenetic distribution of meristic characters mapped onto the BEAST-derived species tree (Clade S = blue, Clade N = orange). (B) Schematic representation of eight diagnostic meristic traits (coded as binary states: red = higher values, blue = lower values). (C) Results of Fisher’s exact tests comparing meristic character states between N and S clade, with p-values indicating significance of inter-clade differences.

**Table 2.**
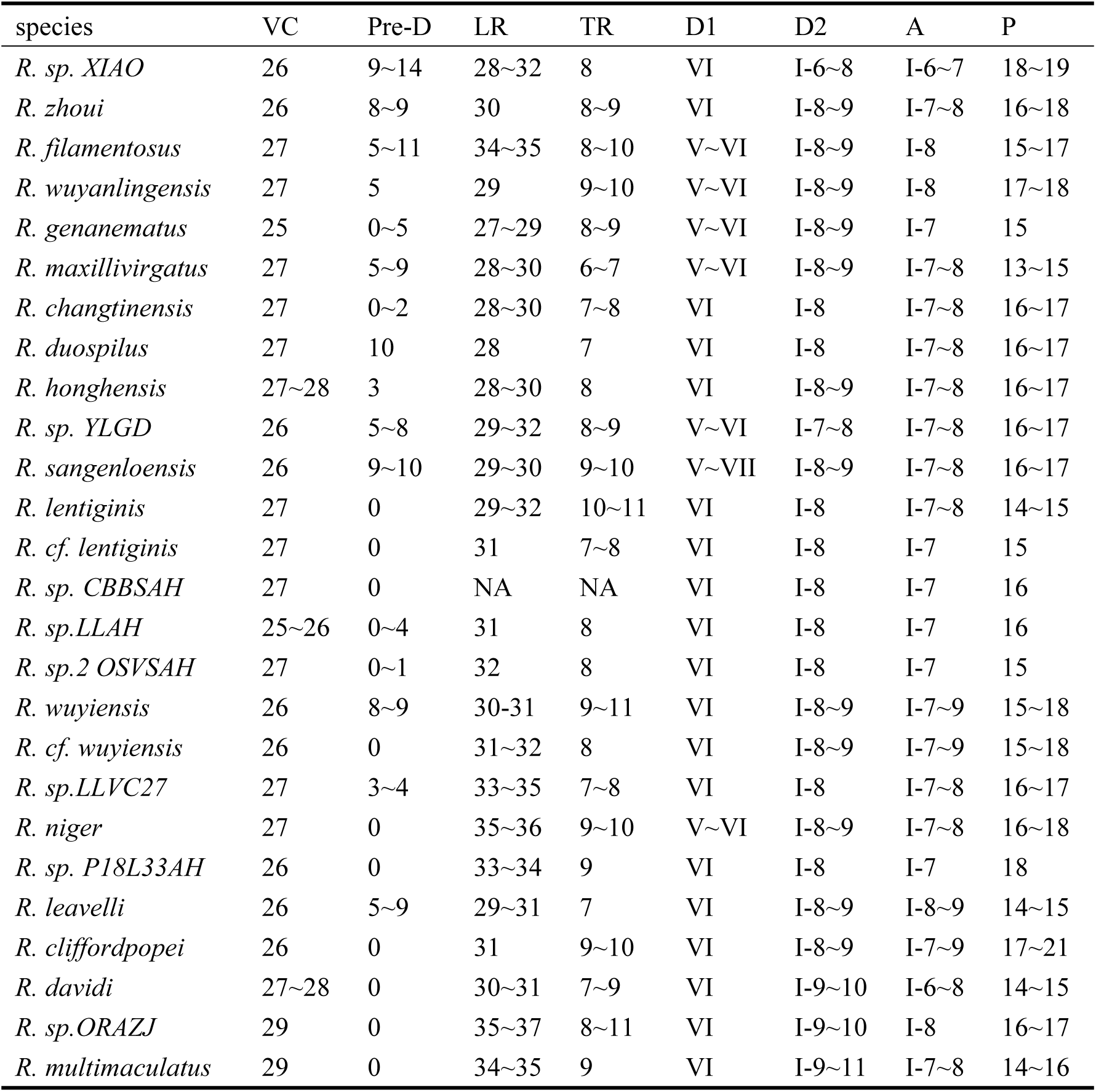
Meristic characters of 26 species of *Rhinogobius* species.

Notably, non-topotypic specimens, e.g., locally collected *R. wuyanlingensis* also showed deviations: pre-D (5 vs 7-9) and lateral scale rows (LR) (29 vs 30-32) (Huang et al. 2016). Historical literature corroborates meristic instability, causing inconsistent species descriptions. For *R. duospilus*, pre-D ranges conflict (8-12 in Wu et al. 2008 vs 6-10 in Li & Zhong 2009), while LR counts vary (29-31 vs 30-32). These findings question the primacy of meristic traits in *Rhinogobius* taxonomy, their variability challenges their sufficiency as standalone diagnostic characters. This concern is amplified by recent species descriptions relying solely on morphological data for affinity assessments (Chen et al. 2024a, b).

#### 3.3.2 Species diversity revealed by phylogenetic, morphological, and geological evidence

Our analysis also reveals that the similarity/proximate among phylogeny, morphology and geological distribution are not always consistent (Fig 3 A). Some sister lineages are geologically close but share few overlapping characters. Specifically in clade-S, *R.* sp. XIAO and *R. zhoui* formed a sister lineage. *Rhinogobius* sp. XIAO processes relatively less second dorsal rays (I-6∼8 vs. I-8∼9 in *R. zhoui*), more pectoral rays (18∼19 vs. 16∼18), and larger predorsal scale variability (9∼14 vs. 10∼12). Notably, these two species are distributed closely (*R.* sp. XIAO in Jiexi county vs *R. zhoui* in Haifeng county, both in eastern Guangdong province), but their coloration is significantly different. *Rhinogobius* sp. XIAO have no organized patch on body side but their upper snout always have an orange stripe whereas *R. zhoui* have 6∼8 big organized orange patch on lateral side and their snout colored white.

Some geological closely distributed species share similar coloration, but they are different in meristic characters and phylogenetically distant. *Rhinogobius duospilus* (typically 27 vertebrae, predorsal scales = 10, first dorsal fin = VI) and *R.* sp. YLGD (26 vertebrae, predorsal scales 5∼8, first dorsal fin = V occasionally) are phylogenetically disparate, but they distributed in two closely located rivers (*R. duospilus* in Rongjiang river vs *R.* sp. YLGD in Hanjiang river of Jieyang City). Conversely, *Rhinogobius duospilus* is clustered with *R. honghensis* (red river basin), and *R.* sp. YLGD is clustered with *R. sangenloensis* (Hainan island), both pairs are distant geographically. Although these clusters consistent with meristic character differences, *R. sp.* YLGD shares most coloration similarity with *R. duospilus*.

The recognition of the *R. lentiginis* complex is further substantiated by a cohesive monophyletic grouping within Clade-N, where *R. lentiginis*, *R.* cf. *lentiginis*, *R.* sp. CBBSAH and *R.* sp. 2 OSVSAH form a well-supported clade. Despite divergent transversal scale counts between sister taxa *R. lentiginis* (10∼11) and *R.* cf. *lentiginis* (7∼8), this assemblage shares synapomorphic traits including near-identical coloration and overlapping meristic ranges. However, *Rhinogobius* sp. LLAH phylogenetically nested in the *R. lentiginis* complex shares less similarities with other species in this group.

#### 3.3.3 Predorsal scales and first dorsal fin spines indicate phylogenetic separation between the southern and northern clades

Furthermore, we assessed the phylogenetic clustering patterns of meristic traits across the phylogeny (Fig. 4B). Predorsal scales exhibited significant phylogenetic signal: species in Clade-N consistently displayed fewer predorsal scales or complete loss of this feature. Absence of predorsal scales was fixed in *R. lentiginis*, *R.* cf. *lentiginis*, *R.* sp. CBBSAH, *R.* sp. P18L33AH, *R.* sp. ORAZJ, and *R. multimaculatus*, but polymorphic (variably present/absent) in *R.* sp. LLAH, *R. wuyiensis*, *R.* cf. *wuyiensis*, *R. niger*, *R. cliffordpopei*, and *R. davidi*. An F-test confirmed significant divergence in predorsal scale absence between the two clades (p < 0.01), These data indicate that predorsal scale loss represents an ancestral state of Clade N, which is supported by ancestors reconstruction result (Supplementary Fig. 2).

Additionally, the stability of first dorsal fin spine counts differed between clades. Species in Clade-N exhibited relatively conserved counts (typically VI), with only *R. niger* showing variability (V–VI). In contrast, multiple species in Clade-S including *R. filamentosus* (V–VI), *R. wuyanlingensis* (V– VI), *R. genanematus* (V– VI), *R. maxillivirgatus* (V – VI), *R.* sp. YLGD(V – VI), and *R. sangenloensis* (V – VII) displayed variable first dorsal fin spine configurations (F-test: p < 0.01). Our result support predorsal scale and first dorsal fin as key meristic characters for species classification.

### 3.4 Reversible predorsal scale loss and recent dorsal fin spine diversification in Rhinogobius

Variation in the number of scales and fin spines is a common feature of adaptive evolution in fishes. Throughout the evolutionary history of ray-finned fishes, scales have been lost on multiple occasions, but once lost, they are rarely regained, suggesting evolutionary irreversibility (Lemopoulos & Montoya-Burgos, 2021). Strikingly, our ancestral state reconstruction of predorsal scales uncovers an exception: this trait was lost in the common ancestor of the northern *Rhinogobius* clade but subsequently re-evolved in *R. wuyiensis*, R. *sp. LLVC27*, and *R. leavelli*. (Supplementary figure 2).

Dorsal-fin spines, another adaptive trait linked to predator defense (Price et al., 2015), exhibit marked divergence between the northern and southern *Rhinogobius* clades, similar to landlocked sticklebacks (*Gasterosteus aculeatus*), which rapidly evolve spine number variations post-isolation (Bell and Foster, 1994). Our study also reveals spine count variability in recently diverged *Rhinogobius* lineages (Supplementary Fig. 2). Evidence suggests that the evolution of dorsal-fin spine number in sticklebacks resulted from the evolution of cis-regulatory sequences in Hox genes (Wucherpfennig et al., 2022). The difference in dorsal-fin spine number between the northern and southern lineages of gobies is likely also driven by similar underlying genetic mechanisms, and selective factors such as convergent evolution under predation pressure change. Collectively, these findings highlight *Rhinogobius* as a promising model for studying evolutionary reversals, adaptive trait diversification, and the genetic basis of morphological evolution in freshwater fishes.

## 4. Conclusion

This study presents the first comprehensive nuclear phylogenomic framework for *Rhinogobius* in mainland China, resolving two deeply divergent clades that originated during the late Miocene (∼9.33 Ma) and subsequent major diversification in the Pliocene. By integrating 2,137 nuclear loci with morphological and biogeographic data, we demonstrate how paleo-drainage reorganizations influenced current distribution patterns, while revealing both the value and constraints of traditional meristic characters for species delineation. Our robust molecular phylogeny provides a critical foundation for future taxonomic revisions, particularly for morphologically cryptic lineages within *Rhinogobius*. The pronounced north-south divergence patterns and dynamic trait evolution uncovered here highlight the need for further genomic studies to elucidate the mechanisms of freshwater adaptation. We recommend expanding sampling to underrepresented regions and employing population genomic approaches to test hypotheses about dispersal routes and speciation drivers. Beyond systematics, this framework offers essential insights for conserving China’s endemic freshwater biodiversity amid growing anthropogenic threats. Ultimately, our findings underscore the importance of combining molecular and morphological data to decipher complex evolutionary histories in rapidly radiating taxa.

## Credit authorship contribution statement

**Suhan Liu:** Conceptualization, Investigation (sample collection, molecular and morphological data), Formal analysis, Writing -original draft. **Qinwen Xu:** Investigation (sample collection, molecular data collection), Validation, Writing -review & editing. **Yun Hu:** Investigation (molecular data collection), Methodology, Writing -review & editing. **Jiantao Hu:** Data curation, Software, Formal analysis, Visualization, Writing -review & editing. **Yiling Pan**: Investigation (morphological data), Formal analysis, Writing -review & editing. **Yun Bu**: Investigation (sample collection), Formal analysis, Writing -review & editing. **Jiliang Wang**: Conceptualization, Resources, Writing -review & editing. **Jianhong Xia:** Conceptualization, Investigation (morphological analysis), Resources, Writing -review & editing. **Chenhong Li:** Conceptualization, Funding acquisition, Project administration, Resources, Supervision, Writing -review & editing.

## Ethical statement

All experimental procedures involving animal specimens were conducted in strict compliance with the Ethical Guidelines for Animal Research established by Shanghai Ocean University (2020). The study protocol was reviewed and approved by the Shanghai Ocean University Animal Ethics Committee (Protocol No. SHOU-DW-2025-019).

## Data availability

Sequence alignment can be found at Mendeley Data, V2, doi: 10.17632/y7dkpb8cs7.2. Raw sequence reads are available from the NCBI Sequence Read Archive (SRA), under PRJNA1311246. All other types of the data that support the findings of this study are available in the main text and supplementary files.

## Acknowledgement

We sincerely express our gratitude to Mr. Huiwen Xiao for help in sample collection. Mr. Zhiwen Huang (Tsinghua University) for his help in GIS data visualization.

## Appendix A. Supplementary

Supplementary materials associated with this article can be found in the online version of this paper.

