## Supplementary figures and images for "Phylogenomics of *Rhinogobius* gobies reveal northern-southern divergence and trait evolution in China"

### supplementary figure.1

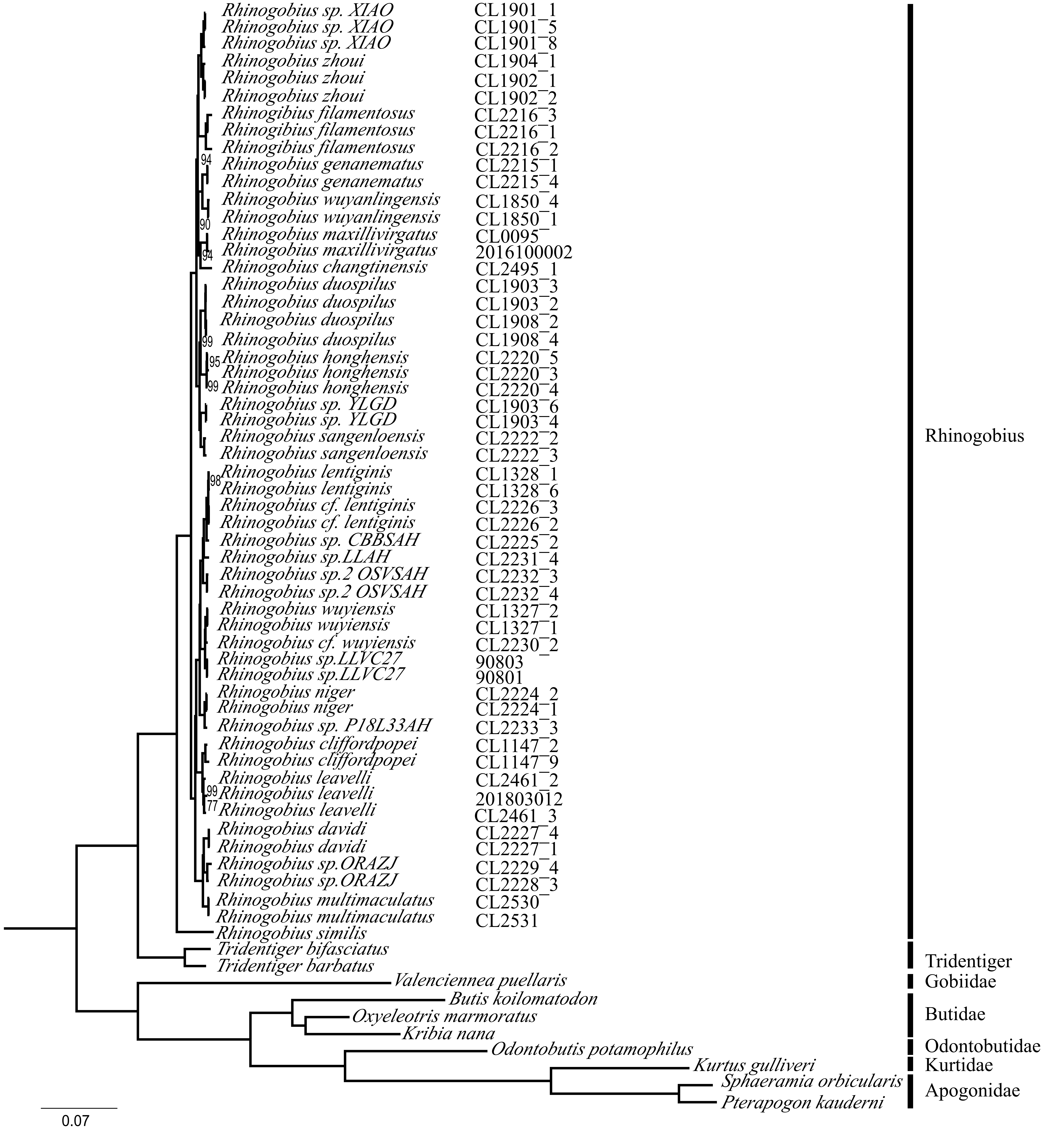

### supplementary figure.2

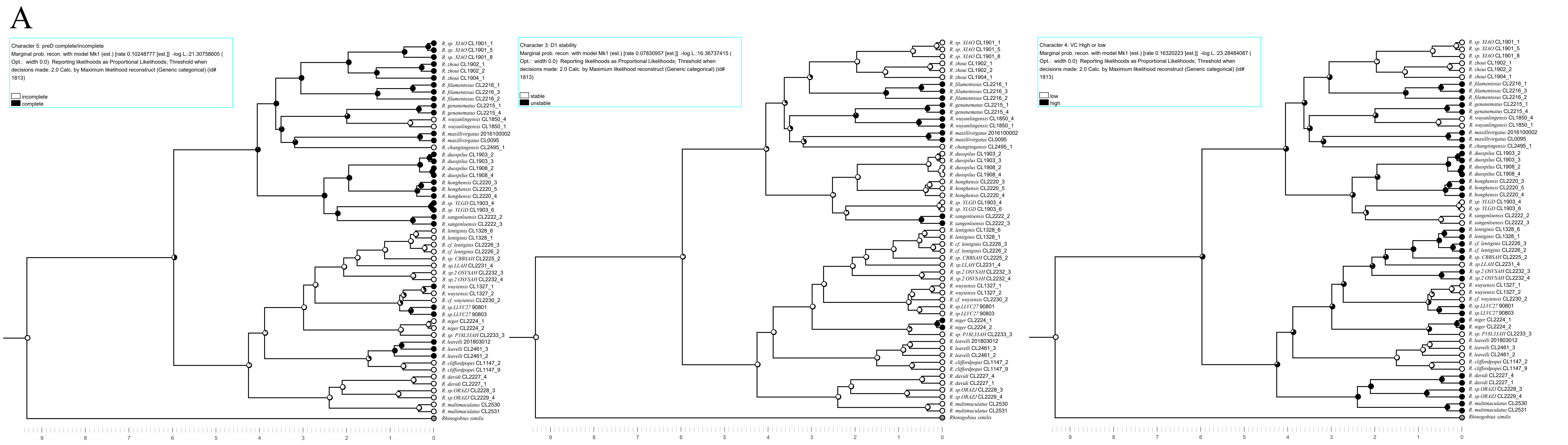

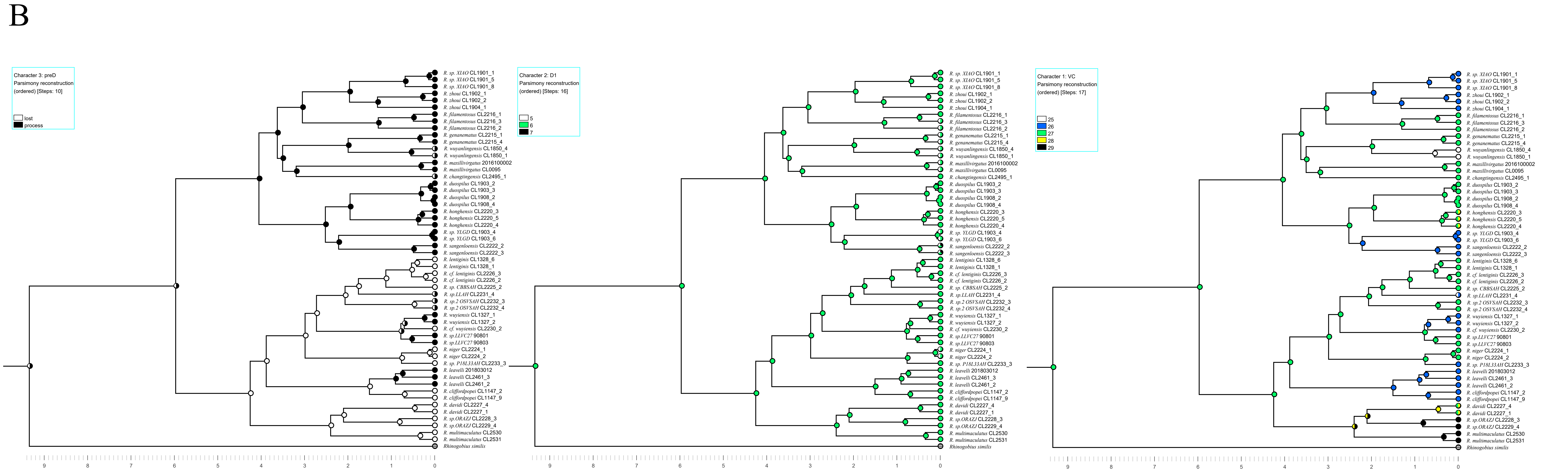
